# Making BrainWaves: Portable Brain Technology in Biology Education

**DOI:** 10.1101/2021.07.02.450935

**Authors:** Ido Davidesco, Steven Azeka, Steven Carter, Emma Laurent, Henry Valk, Jimmy Naipaul, Suzanne Dikker, Wendy A Suzuki

## Abstract

Neuroscience is one of the fastest growing STEM fields, yet its presence in K-12 science education is very limited, partially due to the lack of accessible teaching materials and tools. Here, we describe a new high school neuroscience curriculum (“BrainWaves”), where students utilize recent advances in low-cost, portable Electroencephalography (EEG) technology to investigate their own brain activity. This semester-long curriculum is supported by science mentors and an educational application that guides students through the process of recording and analyzing their own brain activity. Evaluation data collected in 5 public New York City schools in 2018/19 indicate significant positive shifts in content knowledge and self-efficacy among students who participated in *BrainWaves* compared to students in other courses of the same teacher. The quantitative findings are supported by interviews, where students reported increased appreciation for neuroscience and college readiness and discussed the benefits of collaborating with scientists and using portable brain technology in classrooms.

## Introduction

Despite efforts to promote inquiry-based learning, many K-12 science classrooms still rely heavily on the direct transmission of pre-packaged knowledge and overly orchestrated practical work, ultimately restricting students from experiencing the open-ended investigative nature of science [1, 2]. This raises questions with regard to the effectiveness of existing practices: Science learned in highly structured settings may not effectively generalize beyond the school walls to real-life contexts [3, 4]. Therefore, there is a critical need in STEM education for inclusive, classroom-based research experiences, especially at the high school level.

Here, we describe an innovative high school curriculum that provides students with classroom-based research experiences related to cognitive neuroscience. Neuroscience is one of the most rapidly growing STEM fields with increasing attention from public media [5]. Yet, the presence of neuroscience in K-12 education is limited, and students and teachers are often unaware of the field’s foundations and dramatic advances [6, 7]. A larger presence of neuroscience content and access to the scientific process in the high school curriculum could have a number of positive effects on students’ academic life. For example, it enables students to critically evaluate the (neuro)scientific research presented by public media [8]. By exposing students to brain research on the consequences of lifestyle choices, such as the negative effects of drug abuse and positive effects of physical exercise, teachers can encourage students to adopt a healthy lifestyle [9,10]. Importantly, neuroscience can be very easily related to students’ everyday life (how do we learn? how can we improve our memory? how does music affect our brains?). Thus, neuroscience research can provide a gateway into engaging young minds in science and significantly broaden participation in STEM via students’ curiosity about themselves and their peers. Related to this point, neuroscience is also unique in its interdisciplinary nature: By studying the brain, students observe how principles of physics, biology, chemistry, math and computer science are all implemented to gain an understanding of a complex biological system. Finally, introducing students to the concept of brain plasticity can transform the way they think about their learning process and possibly influence their growth mindset, i.e. their belief that intelligence can be developed [11].

A larger presence of brain research in science education aligns well with the Next Generation Science Standards (NGSS) [12]. In the process of learning about the brain, students can internalize crosscutting concepts that apply to different science domains (e.g. the connection between structure and function; division of labor: different parts of a system have different functions). Moreover, several disciplinary core ideas, such as information processing, are tightly related to neuroscience. For example, according to the NGSS, by the end of grade 12 students are expected to understand that “the brain is divided into several distinct regions and circuits, each of which primarily serves dedicated functions” and that “the integrated functioning of all parts of the brain is important for successful interpretation of inputs and generation of behaviors” (p. 150) [12].

Teaching neuroscience in schools poses a few challenges. Teachers typically do not have the background knowledge and tools to teach neuroscience effectively. Research shows that educators (as well as students) often harbor misconceptions about the brain (“neuromyths”), such as “we only use 10% of our brain power” or “there are left- and right-brain learners.” In many cases, such neuromyths are used to justify ineffective or unassessed approaches to science teaching [13, 14]. Furthermore, the price tag of neuroscience research equipment (e.g. brain scanners) and the expertise needed to operate it make it impractical to bring it to schools.

Here, we describe an innovative approach to STEM education in which high school students become brain scientists in an original study of their own creation. Students in the “BrainWaves” program are provided with the content knowledge and practices to design and conduct a comprehensive neuroscience research study in their own classroom with the use of low-cost portable brainwave measuring devices (electroencephalography, or EEG, headsets). Our classroom-based approach supports inclusive implementation. Rather than taking students out of school and into a science laboratory, we bring the laboratory (including science mentors) into the school. Through collaboration with a neuroscientist, students generate a research question, form a hypothesis, design an experiment, collect and analyze brain data and finally present their findings.

## Methods

### Curriculum Description

*BrainWaves* is a semester-long neuroscience curriculum where students conduct hands-on labs and develop their own neuroscience experiment in small teams. The curriculum follows the 5E instructional model [15] and has been developed using backwards design in alignment with the Next Generation Science Standards (NGSS) [12] and the Understanding by Design framework [16].

The curriculum consisted of two units (see Fig. 1). The first unit focused on cognitive neuroscience foundations and included three distinct parts: (1) *How do neurons work?*, (2) *How do different parts of our nervous system work together?*, and (3) *How does the brain interact with the environment?*. In Part 1, students began by exploring the function of neurons by using a bioamplifier called a SpikerBox [17]. Through this lab, students learned about action potentials by recording electrical impulses from insects. Students also created models of neurons and reviewed articles on the function of neurons. Part 2 focused on brain structures through a sheep brain dissection. Students also investigated the functions of each brain lobe through case studies. In the final part, students investigated the concept of brain plasticity through a mirror tracing experiment where they had to repeatedly trace a star by looking at reflection of their hand in a mirror and measure their rapid improvement in the task.

**Figure 1.**
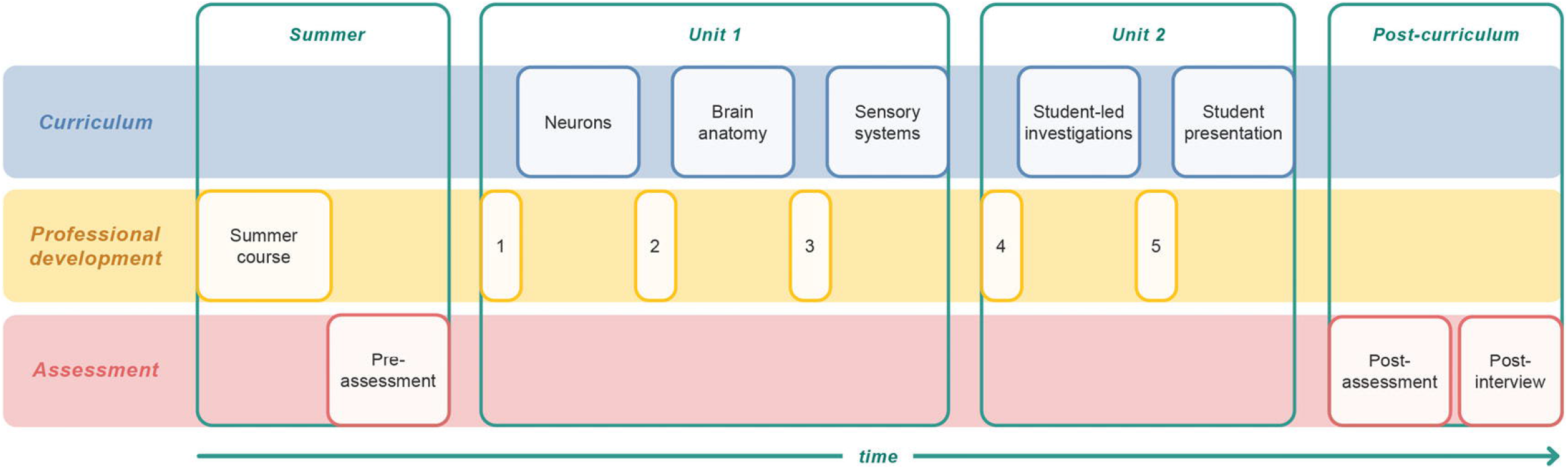
Program timeline. The program began with a professional development (PD) summer course for teachers and mentors with additional PD throughout the implementation of the program. The curriculum spanned over a semester and consisted of two units. Program evaluation consisted of pre- and post-program surveys as well as semi-structured interviews.

At the conclusion of each part of Unit 1, students were introduced to a different aspect of the scientific method. For example, following the SpikerBox experiment students learned about developing research questions and conducting background research. In the following part, students developed hypotheses, began to identify independent and dependent variables, and learned about study design. Lastly, students gained experience collecting and analyzing data through the mirror tracing experiment. By applying the scientific method to each part of the Unit 1, students were prepared for Unit 2 where they would develop their own experiments.

Unit 2 centered on students working in small groups to create their own neuroscience experiments. With support from their teacher and science mentor, students completed all the steps researchers would typically conduct in a research study. They began by developing their own research questions. Combining the content learned in Unit 1 and their personal interests in topics such as exercise, gender differences, and music, students with similar interests formed groups and started developing a research proposal. Once proposals were reviewed by mentors and teachers students began data collection and analysis. Many students’ experiments utilized EEG headsets and the *BrainWaves* computer application, which were two new forms of technology for students. The program culminated with the *BrainWaves* symposium, where students shared their findings with program participants from all the schools.

### Professional development and program implementation

In the summer of 2018, eight public school New York City teachers participated in a five-day professional development course (with science mentors joining them on the third day). The course covered the structure and function of neurons, brain anatomy, and the neural basis of sensory systems. It also introduced teachers and mentors to labs and to the process of conducting research with students. Teachers and mentors also attended five additional professional development sessions during the implementing semester to review key labs and address any programmatic needs (see Fig. 1).

All of the schools participating in the *BrainWaves* program implemented the program as a science elective course that met daily during regular school hours. Students spent the first half of the semester learning neuroscience content and the second half conducting their own original neuroscience research (see Fig. 1). Each participating school was paired with a mentor, typically a doctoral or postdoctoral neuroscientist, that visited the school once a week to help support implementation. At the end of each implementing semester, students from participating schools gathered to present the results of their neuroscience research in a culminating symposium.

As an additional support, the *BrainWaves* application was created as a one-stop-shop for high school students to customize and run their own neuroscience experiments. The application allowed students to 1) run through a classic Event Related Potential (ERP) experiment where the brain response to images of faces and houses was investigated, and 2) customize an experiment where students could change the type of stimuli presented and the timing of their presentation. In conjunction with the BrainWaves application, students used portable EEG devices (Emotiv EPOC+) to measure their own brain activity.

### Program Participants

Eight public New York City schools participated in the program in 2018/9. Each participating teacher was asked to identify another science course that they teach to be used as a comparison group. One school did not complete the program due to scheduling conflicts, and two schools that were transfer high schools were also excluded due to challenges with student attendance and obtaining parental consent. Thus the final sample consisted of five schools.

The experimental group consisted of 169 students and the comparison group consisted of 102 students. To be included in final analysis, students had to provide consent and parental consent, attend more than 75% of the lessons, and complete both the pre- and post-program surveys. The final sample consisted of 86 experimental students and 63 comparison students. Detailed demographics for the experimental and comparison groups can be found in Table 1.

**Table 1:**
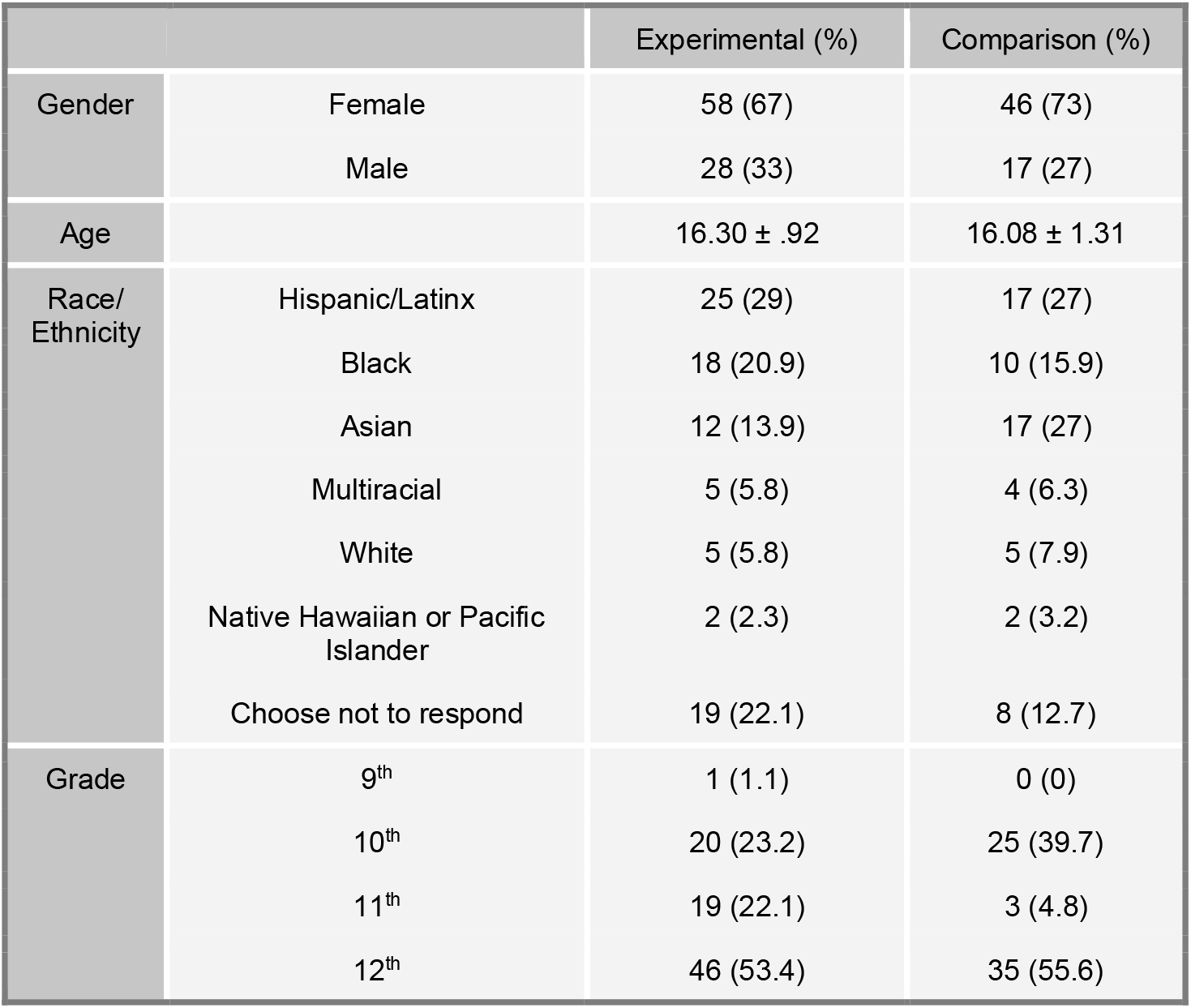
Program participants.

### Program evaluation

A Student Survey was developed and administered to all study participants during the first week of the semester (pre-test) and again during the final week of the semester (post-test). The student survey measured the following constructs:

1. Neuroscience Content Knowledge: building on previous research [13, 14], the survey measured students’ ability to identify misconceptions about the brain (“neuromyths”). Students were presented with 20 statements about the brain and were asked to indicate whether each one of the statements is true or false. Sample phrases are: We only use 10% of our brain power; Circadian rhythms (“body-clock”) shift during adolescence, causing students to be tired during the first lessons of the school day; Some students are right-brain learners, and some are left-brain learners. Half of the statements were true and half were false. These items have been researched among primary and secondary school teachers across various national contexts and adult populations [13]. In addition, students were asked to indicate what percentage of their responses were guesses (none; less than 25%; between 25% and 50%; between 50% and 75%; more than 75%; all).
2. Self-efficacy was measured with a pre-post survey, where students rated their understanding of neuroscience principles (e.g., “my understanding of how neurons communicate with each other”) and research skills (“My skills at using data to understand a research topic”) on a six-point Likert-type response scale (1 = low, 5 = high). In the post-program survey, students were asked to rate their current level of understanding or skill as well as retroactively report their level at the beginning of the program [18]. Neuroscience self-efficacy was only assessed in the experimental (BrainWaves) group.
3. Science interest was measured with the STEM Career Interest Survey [19]: Eleven 5-point Likert-scale (strongly disagree to strongly agree) items assessing students’ self-efficacy, personal goal orientation, outcome expectations, interest, contextual support, and personal input regarding STEM careers (e.g., “If I do well in science classes, it will help me in my future career”).

The assessment process also consisted of semi-structured interviews with teachers and students at the conclusion of the program. Across the five schools, 17 students (9 females) and five teachers (all female) were interviewed for a total of 22 interviews. To reduce bias, all interviews were conducted by two qualitative researchers who were not involved in the development or implementation of the *BrainWaves* program. Interviews were semi-structured; an interview guide was created but researchers were encouraged to ask their own follow-up questions to gain greater content and context from the participants on areas that needed more explanation and depth. All of the interviews were conducted in the schools after consent was obtained. The interviews were audio recorded and later transcribed.

The analysis of the interview was completed by a qualitative research team, which consisted of two researchers who contributed to the development and implementation of the BrainWaves program, and the two interviewers. The process of horizonalization was used to code the transcripts, which is when every statement is analyzed equally and significant statements that provided understanding of the phenomenon were coded and highlighted [20] The significant statements were used to generate larger clusters of meanings, or codes. The two interviewers then re-analyzed all 22 transcripts using the 18 codes that were established based on quotations from teachers and students to assure that the themes were grounded in the data. Codes were then grouped together based on commonalities to determine overarching themes. These themes were then checked against the transcripts to assure that they were representative of the participants experiences.

## Results

### Survey data

We first assessed the potential impact of the *BrainWaves* program on students’ self-efficacy in neuroscience. Paired t-tests revealed significant gains in students’ self-efficacy in neuroscience from pre (mean = 2.232; SD = .872) to post (current) (mean = 3.868; SD = .740; *t*(85) = 15.957, *p* < .001) and from retrospective (mean = 3.031; SD = 1.122) to current (mean = 3.868; SD = .740; *t*(85) = 6.430, *p* < .001) (Fig. 2A).

**Figure 2.**
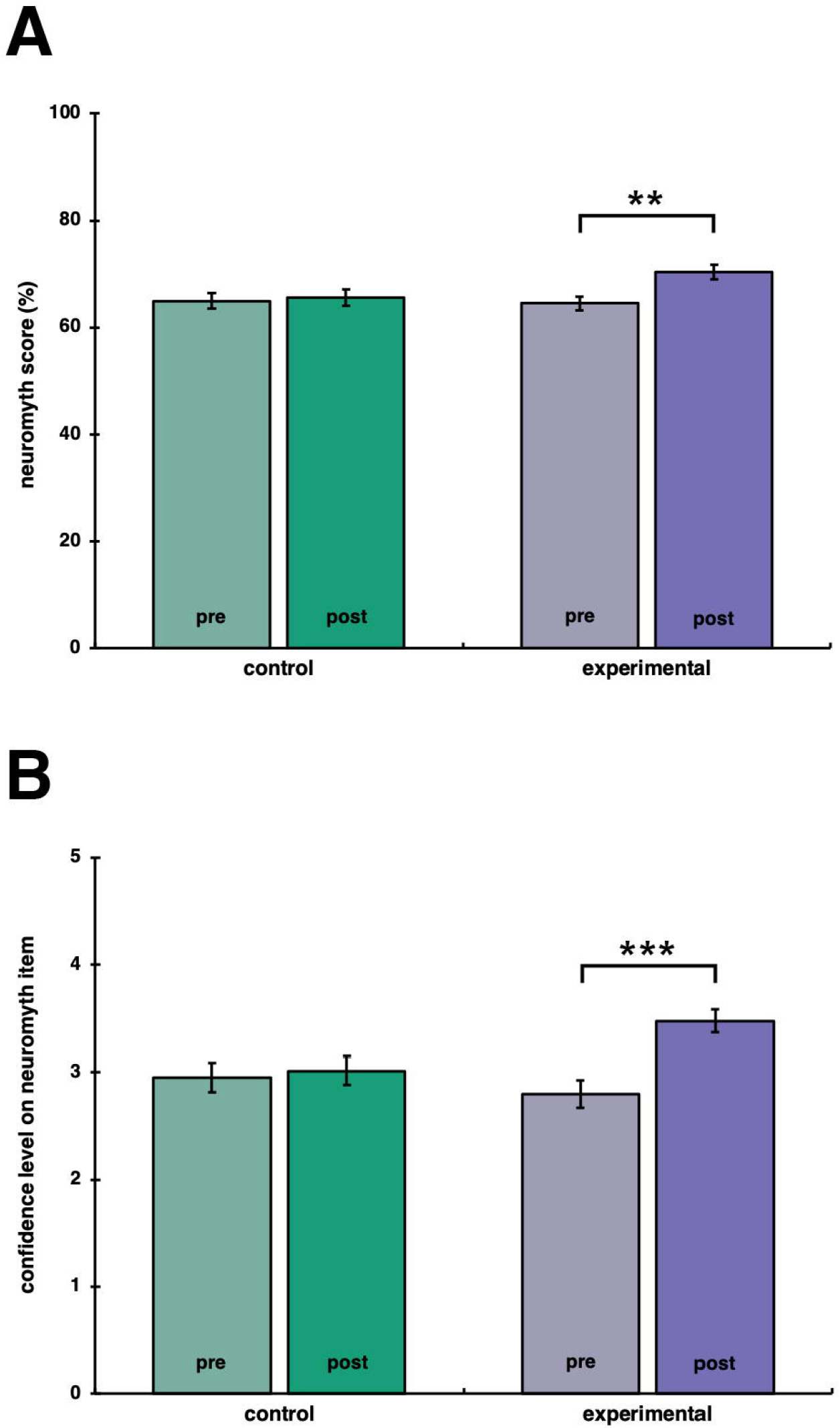
Self-efficacy in neuroscience and research. (A) Self-efficacy in understanding of neuroscience principles before and after the program. (B) Self-efficacy in research before and after the program. In the post-program survey, in addition to rating the current level of skill or understating, students were asked to retroactively report their self-efficacy at the beginning of the program (striped bar). Colored bars depict mean score; black bars represent standard error of the mean. ** p ≤ 0.05. *** p ≤ 0.005.

A similar effect was observed in students’ confidence in their ability to identify misconception about the brain (“neuromyths”). A two-way mixed ANOVA revealed a significant time by group interaction (F(1, 147) = 7.160, p = .008, n_p_^2^ = .046), with BrainWaves participants showing larger increases in their confidence from pre (mean = 2.791; SD = 1.189) to post (mean = 3.453; SD = 1.013) relative to the comparison group (pre: mean = 3.034; SD = 1.042; post: mean = 3.052; SD = 1.083) (Fig. 3B). However, there was only a non-significant trend in students’ actual content knowledge scores (F(1, 147) = 3.504, p = .063, np2 = .023) (Fig. 3A).

**Figure 3.**
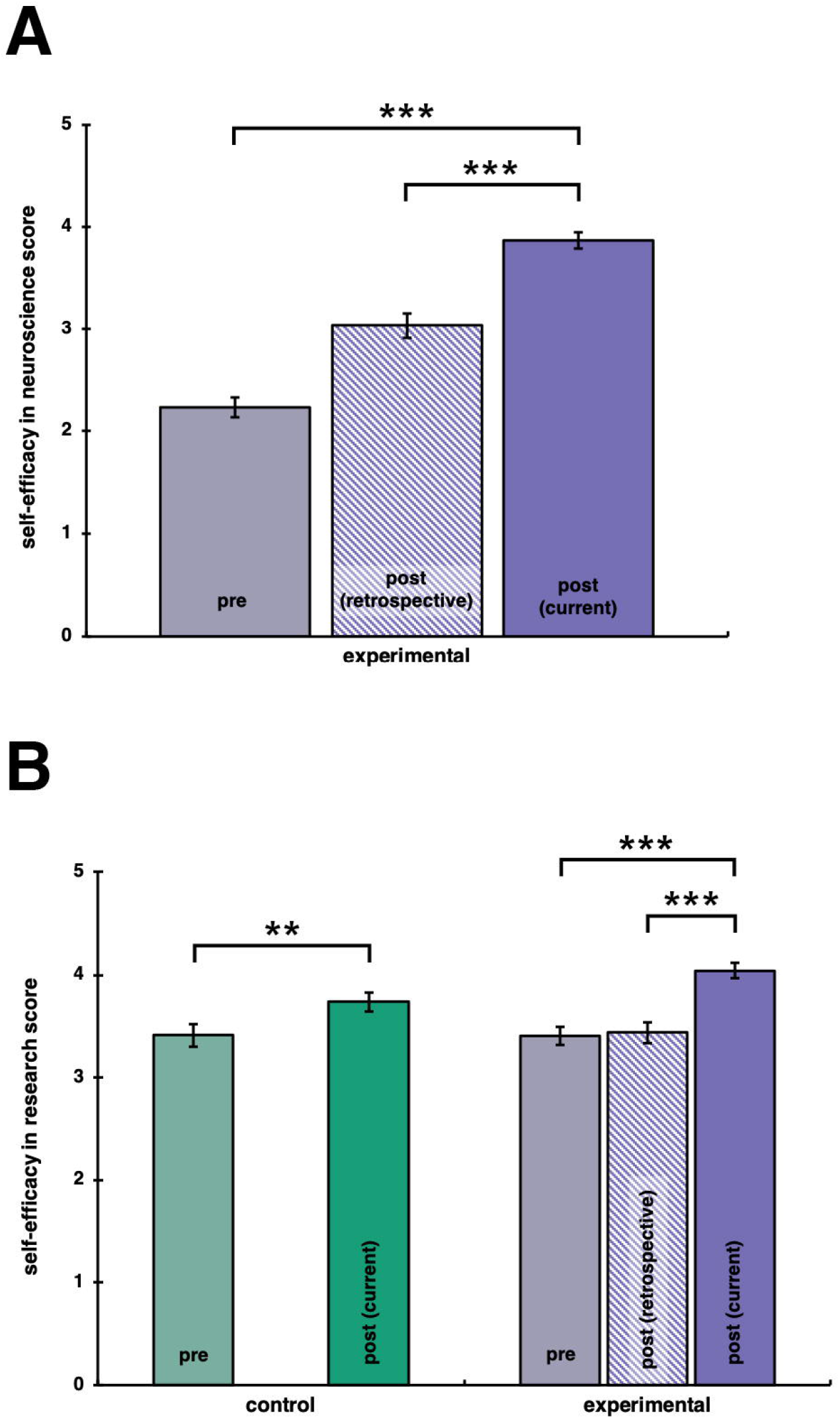
Students’ content knowledge. (A) Proportion of correct answers in pre and post-program knowledge tests; (B) Students’ confidence level in their answers.

Next, we assessed the potential impact of the program on students’ self-efficacy in research. A two-way mixed ANOVA showed a significant time by condition interaction (F(1, 147) = 4.337, p = .039, n_p_^2^ = .029), with BrainWaves participants demonstrating stronger positive shifts from pre (mean = 3.41; SD = .806) to post (mean = 4.044; SD = .707) relative to comparison participants (pre: mean = 3.413; SD = .894; post: mean = 3.738; SD = .744) (Fig. 2B). Despite the significant increases in students’ self-efficacy and knowledge confidence, there was no significant time by group interaction for students’ science interest (F(1, 147) = 0.981, p = .324, n_p_^2^ = .007) (Fig. S1).

### Semi-structured interviews

Four themes emerged from the analysis of student and teacher interviews:

#### (1) Appreciation for Neuroscience

##### Students

Several students acknowledged the uniqueness of the BrainWaves program stating, “…science is usually memorization and like practice makes perfect so neuroscience was a lot more hands on and fun than other science classes so it definitely changed my perception” (17-year-old male student). Another student stated how the program led to more interest: “So we got to learn a lot more than just looking in a paper and just reading it. You are a lot more engaged. So I think it does help students get more interested into science” (16-year-old male student). The authenticity of conducting neuroscience research was also appreciated by students:

> *I liked it because it is different from what I am used to working with and it feels more real because like those EEGs they were like I don’t know it makes you feel more like you are a scientist or something it makes it feel more serious and it really looks cool and stuff like you really get signals of the brain and it is just real. (16-year-old female student)*

Other students reflected on the impact of the program on their perception of health and wellbeing: “It made me more careful with what I do with my body because my brain is everything so I have to be careful with what I do” (17-year-old male student). Another student agreed, adding that, “…I feel like neuroscience relates to you and like I said, I feel like when something you’re learning about relates to you you’re more likely to listen… you’re more likely to want to learn about it.” (16-year-old female student). Another student commented on how the curriculum would be beneficial for his future stating, “A lot of people I know want to learn more about themselves and like their brains and everything, and also careers that involve brains. It is an important system and I think it is important to understand more” (17-year-old female student).

##### Teachers

Similarly, teachers described an overall impact on students’ thinking noting that the program has “…prompted them to ask questions and to delve deeper… it really does make them think beyond the obvious”. (Female science teacher of 8-years from Manhattan). Teachers also reported that the program impacted their teaching:

> *“It also allowed me to brush up on a lot of my neuroscience knowledge and thinking a lot about bringing the ideas to high school students in a way that they would understand it like it is still complex, um, but they can still process the information that I am teaching*.*”* (Female teacher of 10 years).

#### (2) College Readiness

##### Students

Over 70% of the students in BrainWaves were in their final two years of high school (Table 1), and therefore college admissions were brought up in several student interviews. Students expressed how being part of the BrainWaves program strengthened their college applications. “The other benefit I would say is applying to college and wanting to be a scientist and you tell the school I did a neuroscience class it will be a little better for getting into the school you want to.” (17 year old female student). Students also expressed how the program made them think about what life would be like in college:

> *“I think it prepares us for college because if we have that experience and exposure now we won’t receive that as a shock in college and say, ‘Oh my god, what is this’*… *we need to know now so that we are not overwhelmed at college and end up failing the course*.*”* (16 year old male student)

Additionally, students reported that the independent research projects they completed in the BrainWaves program would be similar to the independent work they would have as college students: “It taught you how to work with that kind of freedom because in college no one is going to tell you what to do. It is your own responsibility to actually do the project” (17 year old male student). Lastly, some students expressed interest in pursuing neuroscience in college: “it has affected my decision making by possibly making me change my major in college…I am considering neuroscience because I really want to learn more about the brain and who knows someday maybe neurosurgery” (17 year old female student).

##### Teachers

The concept of the BrainWaves program leading to a greater sense of college readiness was widely endorsed by the teachers in the program. The teachers attributed this sense of readiness to the students designing and completing an independent research project, which was the focus of Unit 2 of BrainWaves. Teachers believed the research experience gained from the program provided students with the skills necessary for rigorous college courses:

> *“I think is really important and here we are still holding their hands to a certain extent but I think this is a good way for them to see what it will be like in college, um, whether they are presenting or writing papers, I think it was a really good way for them to prepare for college readiness*.*”* (Female teacher of 10 years).

In addition to independence, teachers noted that reading and writing academic reports were unique skills that students would not have been taught as part of a typical high school curriculum. For example, a teacher of 15 years from the Bronx stated, “It was my freshman year in college I think my lab teacher kept giving me C’s and I couldn’t understand why and it was because I didn’t have any experience writing labs in science like zero experience from New York City high schools and to think that is how it was 16 years ago and nothing has changed.” A similar sentiment was expressed by a science teacher of 9-years from the Bronx, who compared the BrainWaves program to her own college research experience:

> “The first time I had to look at a journal article probably wasn’t until sometime later in college and I definitely think I was intimidated, whereas these students are going to know how to walk into the library and figure out what publication they need; they are going to know that even though this looks really dense, they are going to be able to figure out as much of it as they need…”.

#### (3) Support from Mentors

##### Students

Many students discussed the support they received from the neuroscience mentors. The students acknowledged that the mentors were field experts and that it was their expertise that helped them understand and solve issues with their own experiment:

> *“They definitely helped out with collecting the data and then putting it into a chart. They also helped us come up with ideas because most of the class was confused about what type of experiment to do and what it has to do with the brain, so they helped us. They pointed out important parts of the brain and said maybe we could talk about that area*.*”* (17-year-old female student)

The mentors became a fixture in the students classrooms and provided students with the guidance necessary for them to grow. An 18 year old student stated, “They were very helpful and guided us a lot and they always made sure that, um, we could go up to them and ask them questions. They weren’t just an extra person in the room.” The mentors also provided the students with direct exposure to science careers. One student stated of their mentor:

> *“… help people kinda get out of their comfort zone so that they can realize there is a huge market for something. There are opportunities if you have someone from other industries from other lifestyles career paths to come in and say hey we exist, here are some interesting facts like careers learn more about careers*” (17-year-old male student)

##### Teachers

Teachers also reported feeling supported by the neuroscience mentors. Echoing the students’ sentiments that the mentor was more than “just an extra person in the room”, teachers appreciated the expertise the mentors had with the neuroscience content.

> *“They acted as an expert… to sort of answer questions that I couldn’t answer about the content or the process and she would answer them, either the context process or the tech. So she was like a backup expert in the room. It was great”* (Teacher of 6 years)

Several of the teachers acknowledged that they were not trained in neuroscience and that they had to learn much of the content before teaching their classes. The mentor’s expertise with the content and familiarity with the technical components, such as the EEG, was found to be extremely helpful for teachers. As the semester progressed past the content portion, the mentors supported the teachers as the students began developing their own research: “The mentors came in and they really helped me when it came to checking the proposals before they started it and checking their research questions and methods, which was all part of their proposal” (Teacher of 8 years).

Teachers expressed that even if they knew the material or aspect of the research, it resonated differently with the students when delivered from a mentor. A teacher of ten years stated, “I think having someone coming consistently, they are able to build a relationship with them, they are able to ask them questions about the field, and they also respect them more”. The teachers expressed that the mentors were a unique aspect of the BrainWaves program and was something they believed the program should continue to implement as it moves into future iterations.

#### (4) Use of Technology in Science Classes

##### Students

The use of technology in BrainWaves was a key topic present in many interviews. Two forms of technology that were novel to the students were the electroencephalogram (EEG) headset and the BrainWaves software application. One student stated, “We never really used technology in our prior science classes. We never had the opportunity to use EEG headsets. Also, we never had the opportunity to have a class dedicated to neuroscience” (18-year-old male). Another student reported that unlike *BrainWaves*, in other science courses:

> “We didn’t really use much technology we just used textbooks and just needed to write things down and learn formulas and stuff we didn’t really use much technology only sometimes if we had to write something on the computer or had to write some results that was it.” (15-year-old female student)

Students were cognizant that they were not always engaged in their previous science classes. They described how better access to technology through the BrainWaves program improved their attentiveness and engagement in science class. A 16-year-old male student said:

> *“Using technology is fun and I guess it is something that most kids will not have access to or like students will not have access to, to these technologies, to explore with. The amount of possibilities you have are large, so just having fun and experimenting with it, um, looking around and seeing what it can do. All of this was a fun part and that is why I think it was really good for us. It got us interested and it got us motivated to do this work*.*”*

##### Teachers

Teachers also commented on how the technology in the program increased students’ engagement. The EEG device was used when the students were conducting their research experiments and provided a practical link to the concepts they had learned previously in the course:

> “The real time feedback from the brain was fascinating. They could see their brainwaves being picked up and what they have been learning and what I have been saying in class is actually real. And it is not like we are just talking about these electrical impulses but here they are on the screen.” (Teacher of 10 years)

As with many of the concepts discussed in these interviews, technology was also linked to students futures beyond the BrainWaves program. A teacher shared that after having a conversation with a student about how technology, like an EEG headset, can be used in the medical field as well as research, students began to think more about careers in fields that use advanced technology:

> “I think for students to experience real world technology gives them a better idea of what they are looking at when they get out of high school gives them ideas of what they might enjoy and what they might not enjoy.” (Teacher of 13 years)

Although widely regarded as beneficial to the overall program, teachers did address difficulties when using technology. One teacher of 11 years stated, “The technology does engage them but you also need to make sure that they’re engaged in the academic way and that they’re not using a technology to just check social media or play games.”. Additionally, teachers commented on the technical difficulties they faced and how the BrainWaves team and mentors assisted during these experiences..

## Discussion

From the quantitative analysis it was evident that students’ levels of confidence and self-efficacy in neuroscience and in research were positively impacted by their experiences in the BrainWaves program (Figures 2 & 3). The qualitative themes provided two main explanations for why students’ self-efficacy towards neuroscience and research was positively impacted.

First, whereas laboratories in traditional science courses typically follow a cookbook format with little area for students to take ownership over the development and execution of an experiment [1], the BrainWaves program allowed students to authentically design and execute their own research projects.

Second, science mentors provided a sense of comfort for students to ask questions about science topics and careers in research. Similar to the participating students, many mentors came from racial and ethnic groups underrepresented in STEM, providing meaningful role models to students. As indicated in the interviews, by discussing their career trajectory with students, the mentors widened students’ exposure to STEM careers.

### Lesson Learned

The first full year of program implementation, discussed in this paper, proved to have several valuable qualities but is not without its limitations. Our team has identified several challenges associated with implementing the BrainWaves program in public high schools. New York City is the largest district in the United States serving over a million students with a choice - based placement system. As such, students’ prior experiences in science and academic needs vary substantially from school-to-school. During the end of year interviews and in professional development sessions, teachers often commented about needing to modify the provided curriculum to make the scientific language and level of the materials more accessible. Based on this feedback, the BrainWaves team has created several modifications for each lesson to allow for a greater range of implementation.

Another challenge teachers’ faced was pacing. Some teachers reported that activities in Unit 1 took longer than expected. Teachers also reported having technical difficulties with EEG equipment in Unit 2 when students were designing their experiments, which could perhaps explain the lack of change in students’ science interest. With more time to conduct their own research and deeply engage in fewer topics, it is possible that students’ attitudes towards science may improve more in future iterations of the program.

## Acknowledgements

This project was supported by the National Institute of General Medical Sciences, the National Institutes of Health under Award Number R25OD023777. The content is solely the responsibility of the authors and does not necessarily represent the official views of the National Institutes of Health.

