## Supplementary material for "Making BrainWaves: Portable Brain Technology in Biology Education": Figure S1

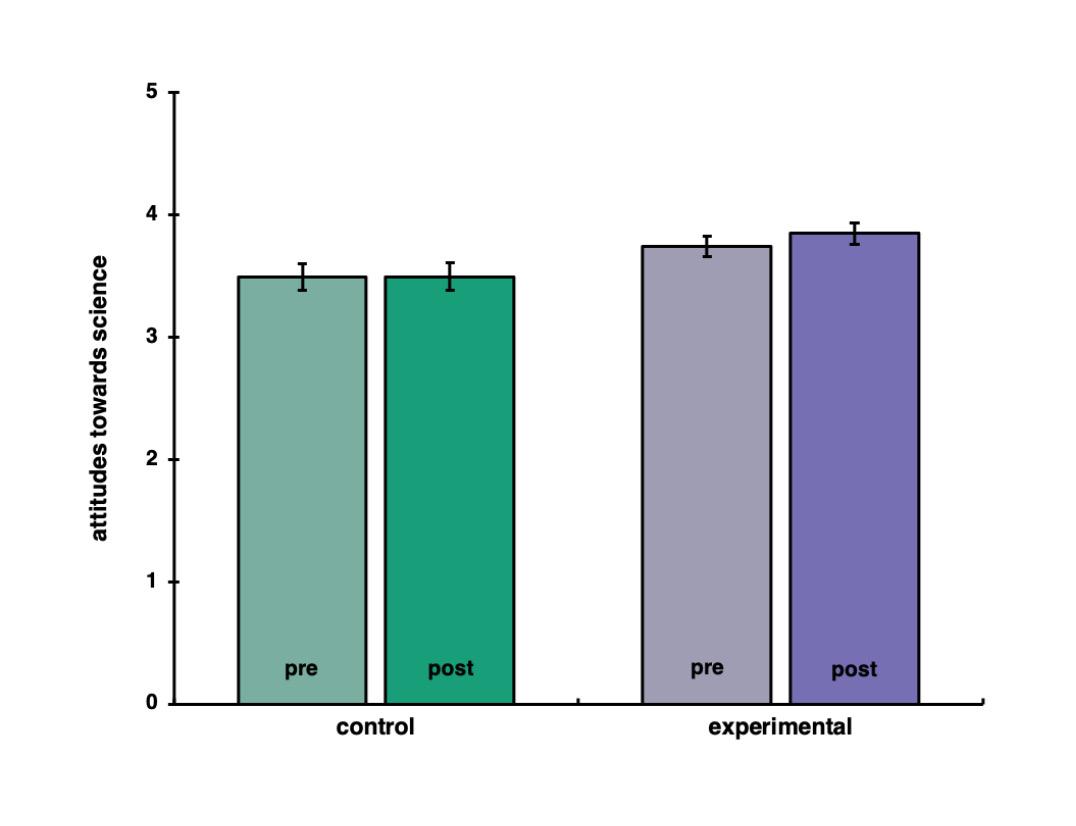

**Figure S1.**  Students’ self-reported science interest before and after the BrainWaves program in comparison to students taught by the same teacher in another science course.
